# High Precision Peptide-MHC Class I CARs Enhance *In Vivo* Killing of Targeted CD8 T Cells to Prevent Type 1 Diabetes in Novel RIP-mOVA Model

**DOI:** 10.1101/2025.11.10.687586

**Authors:** Mathew Prashanth Francis, Winston Hibler, Michael Yarnell, Neetigyata Singh, Ross P Kedl, Matthew A Burchill, Terry Fry, Maki Nakayama

## Abstract

Progress has been made to address refractory T cell-mediated diseases with chimeric antigen receptor (CAR) T cell platforms, with the need more specifically target pathogenic cells being addressed by peptide-MHC (pMHC) CAR T cells. However, progress to date has largely been limited to peptide-MHC Class II (pMHC-II) CARs for the targeting of autoreactive CD4 T cells in multiple models. Here, we develop the first single molecule peptide-MHC Class I-β2 microglobulin CAR platform to demonstrate the proof of concept for depletion of antigen-specific CD8 T cells and compare the *in vitro* and *in vivo* functionality to a traditional scFv-based CAR that also targets the TCR. We demonstrate that while the scFv CAR is more rapidly and robustly activated by target CD8 T cells *in vitro*, the greater precision offered by pMHC-I CARs results in greater efficacy in multiple *in vivo* models. Additionally, we developed a novel adjuvanted ovalbumin vaccination with adoptive transfer model to induce type 1 diabetes (T1D) in RIP-mOVA mice using a polyclonal, polyepitopic T cell population. We demonstrate that our pMHC-I platform depletes SIINFEKL-reactive T cells and that control of the immune response to the dominant SIINFEKL epitope can control a broader antigenic response.

## Introduction

Aberrant T cell activity drives a broad range of diseases including T cell leukemias, solid organ transplant rejection, and many autoimmune diseases – the latter of which affects over 4% of the population.^1^ Historically, many of the T cell targeting therapies for these diseases are characterized by poor ability to discriminate pathogenic from non-pathogenic T cells, resulting in high adverse effect profiles. More recently, anti-CD3 monoclonal antibody (MAb) therapies such as teplizumab have shown clinical success in the prevention of type 1 diabetes (T1D) in high-risk patients, but still suffer from limitations in depth of depletion of pathogenic cells and durability of effect that is seen with many MAb therapies.^2, 3^

Therapies for B cell-mediated diseases share many of these concerns, but recent developments in chimeric antigen receptor (CAR) T cell technology offer treatment for refractory disease. Specifically, anti-CD19 CAR therapy showed remarkable efficacy in preliminary clinical case studies of severe systemic lupus refractory to MAb and immunosuppressive therapies.^4, 5^ As a cellular therapy, CAR T cells offer greater durability of response following single treatment with newer data suggesting the anti-CD19 CAR therapies offer additional advantages over MAbs including broader tissue biodistribution, enabling greater depth of target cell depletion.^6^ However, no corresponding T cell antigens have been found that offer the promise of CD19 targeting for B cell pathologies.

One approach to a greater discrimination and targeting pathogenic T cell is the use of peptide-MHC CARs, which target T cells of interest via binding of their T cell receptor (TCR). This offers the advantage of only depleting T cells that are by definition pathogenic, but requires the presence of immunodominant epitopes for targeting. This overall approach has been used to target autoreactive CD4 T cells in autoimmune models of disease such as autoimmune encephalomyelitis (EAE), autoimmune arthritis, and the non-obese diabetic (NOD) model of T1D.^7-9^

However, to date there are no published peptide-MHC Class I (pMHC-I) CAR formats which target autoreactive, diabetogenic CD8 T cells. Here, we show the development of a pMHC-I CAR within the canonical ovalbumin system. Additionally, we developed an anti-Vβ5 CAR which also targets the most common OVA-reactive CD8 T cells, thus allowing us to compare the kinetics and *in vitro* efficacy of more traditional scFv CARs to peptide-MHC CARs. Finally, we measure their effects against both monoclonal and polyclonal OVA-reactive CD8 T cell populations in established and novel models of autoimmunity.

## Results

### Design and *in vitro* characterization of pMHC-I and anti-Vβ CAR constructs

We designed two CARs to target pathogenic T cells via their TCR. (Fig. 1a) The first is a traditional second-generation CAR with scFv ectodomain which targets the Vβ5 chain of the TCR, which is the β chain found in OT-I CD8 T cells and the largest proportion of SIINFEKL-reactive CD8 T cells. Next, we sought to develop a CAR which would present the cognate peptide and MHC Class I molecule of the TCR. Thus, we developed a second-generation CAR with an ectodomain containing a single chain trimer consisting of β2-microglobulin with the target MHC Class I (H-2K^b^) and immunodominant cognate peptide (SIINFEKL) of the same target CD8 T cells mentioned above. We termed these CARs α-Vβ5 and OVA-H2K^b^, respectively. Because this approach attempts to kill cytotoxic CD8 T cells via binding of their TCR, we opted for a second-generation CAR format biased towards maximizing short term activation with a CD28 co-stimulatory domain and CD3ζ with all ITAM sites able to be activated. Constructs were cloned into pMSCV vector, transfected into Plat-E for retroviral packaging, and then transduced into bulk T cells isolated from mouse spleens. We demonstrated transduction through surface expression of the reporter gene tEGFR. Surface expression of the CAR is demonstrated via anti-FLAG antibody for the α-Vβ5 CAR and anti-H-2K^b^ for the OVA-H2K^b^ CAR. (Fig. 1b)

**Figure 1.**
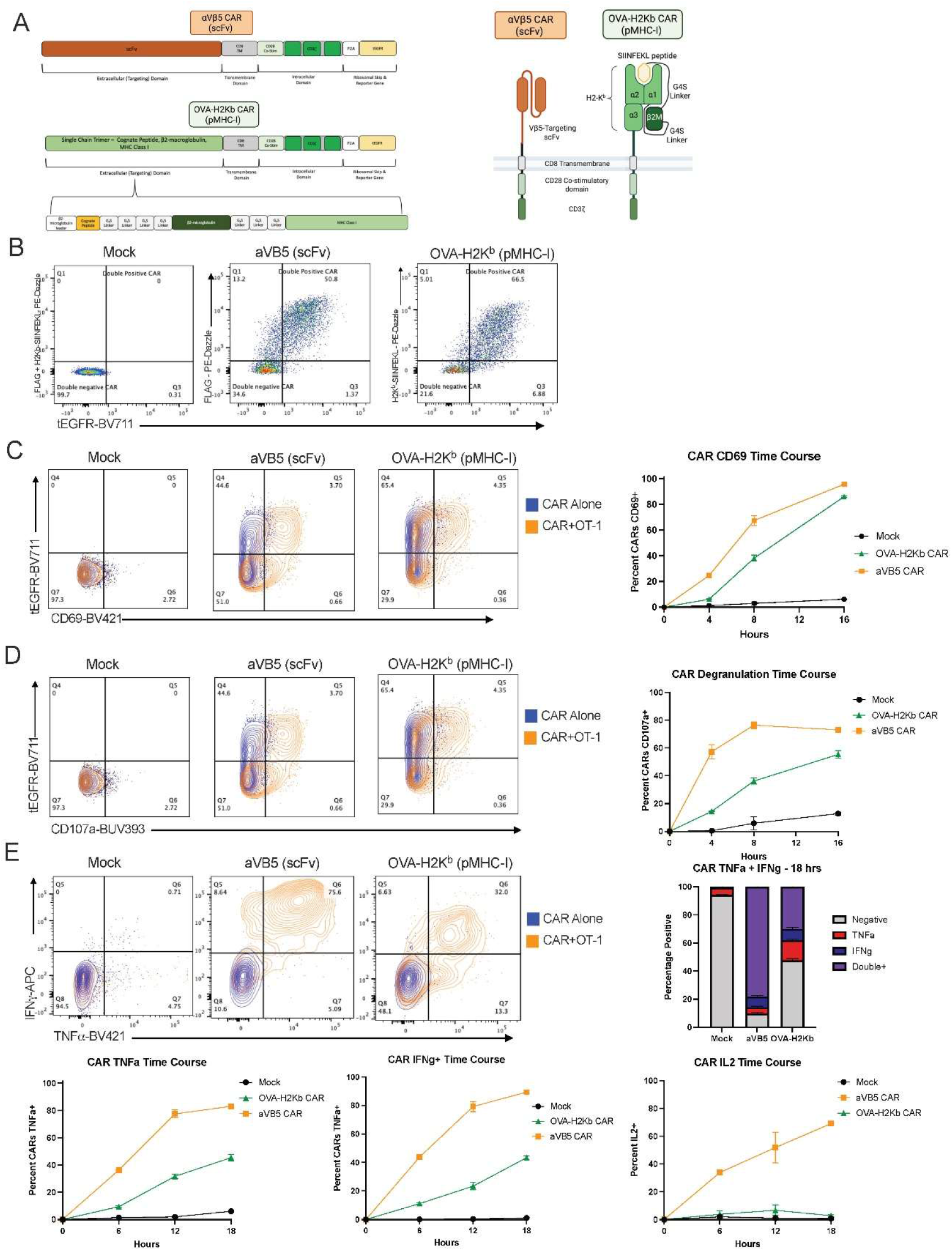
Peptide-MHC-I and scFv CARs display different magnitude and kinetics of activation *in vitro*. (A) Schematic of scFv and pMHC-I constructs. (B) T cells were isolated from naïve splenocytes and transduced with either CAR construct, with good transduction (EGFR) and surface expression (FLAG/H2Kb). (C,D) Co-culture with OT-1 T cells to measure antigen-dependent activation via CD69 and CD107a surface expression as measured by flow cytometry. Time course represents continuous co-culture with OT-1 T cells. (E) Co-culture with OT-1 T cells to measure antigen-dependent intracellular cytokine production. Upper panel and bar graph show dual positive, TNFa+IFNg+ T cells at 18 hr time point, whereas lower panels show time course for single cytokine production of TNFa, IFNg, and IL-2.

To evaluate the functionality of these CARs, co-culture experiments were performed with target OT-I CD8 T cells. Both CARs demonstrated robust, antigen-dependent activation as measured by CD69 and degranulation as measured by surface expression of CD107a, though OVA-H2K^b^ was slower to reach similar thresholds. (Fig. 1c,d). Activation was strongest in CAR T cells which were most effectively transduced, as measured by EGFR expression. While both CARs were able to make tumor necrosis factor-α (TNF-α) and interferon-γ (IFN-γ) in response to antigen, α-Vβ5 CAR T cells displayed a more rapid and robust response and was the only CAR able to produce interleukin-2 (IL-2). (Fig. 1e) This was tested with via intracellular staining and flow cytometry given the co-culture of two T cell populations.

### anti-Vβ CAR T cells kill target OT-I CD8 T cells *in vitro* faster than pMHC-I CAR T cells

Next, we looked at the effects of these CAR T cells on the target OT-I CD8 T cells. As expected, co-culture with the OVA-H2K^b^ CAR resulted in robust activation and degranulation of the OT-I CD8 T cells that was similar to the CARs in magnitude and kinetics.(Fig. 2a) This activation signaling was accompanied by production of TNF-α and IFN-γ, though only minimal IL-2.(Fig. 2b) Interestingly, we observed similar levels of activation from co-culture with the α-Vβ5 CAR despite not presenting the cognate epitope of the TCR. Despite robust activation of the targeted OT-1 T cells, we observed that both CARs killed the target OT-I CD8 T cells.(Fig. 2c)

**Figure 2.**
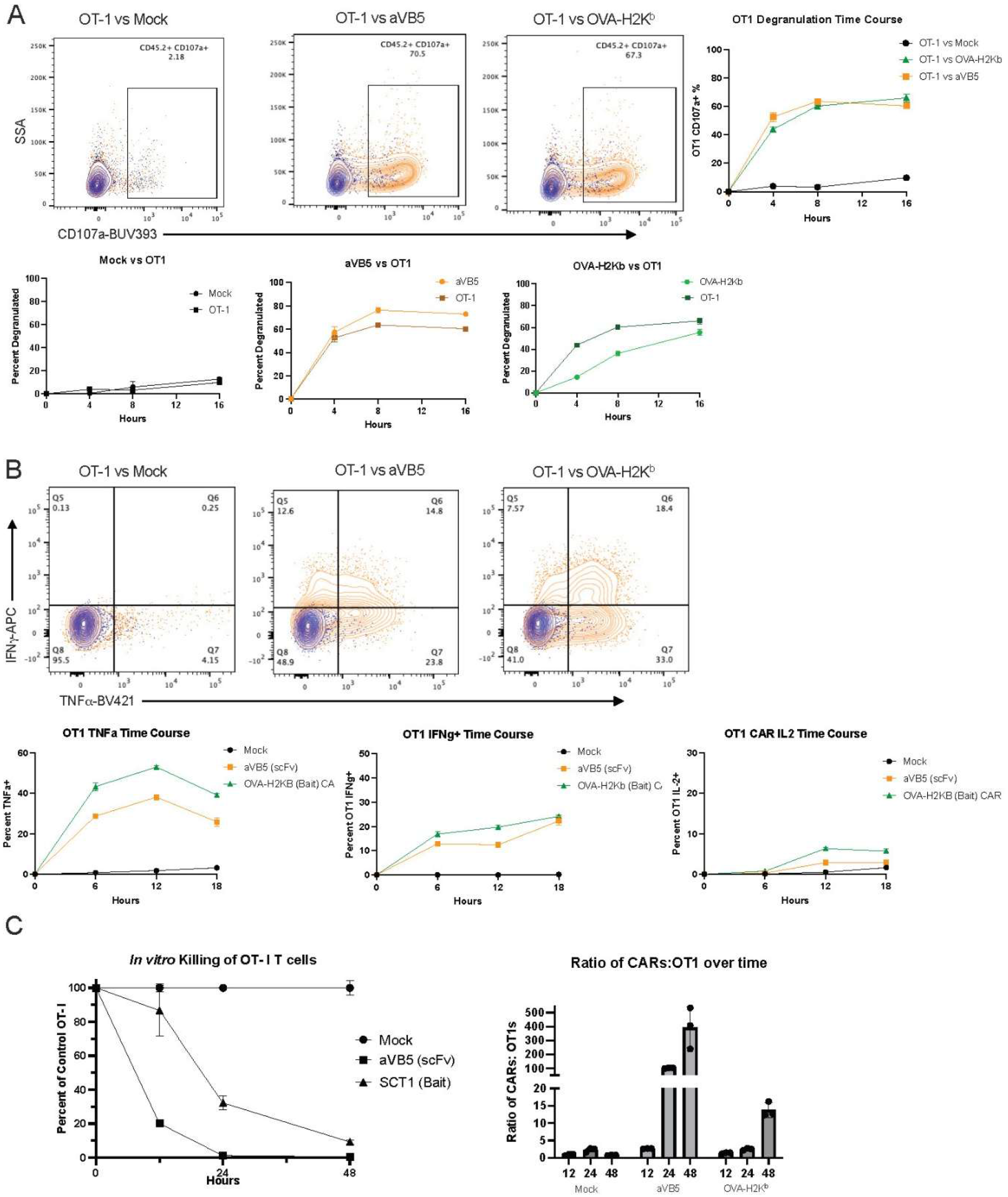
Despite robust activation of target OT-1 T cells, both pMHC-I and scFv CARs kill target OT-1s. (A) Shows only OT-1activation when in co-culture with listed CARs, as measured by CD107a. Right panel shows time course of OT-1 degranulation when in co-culture with listed CAR. Bottom panels compare OT-1 degranulation kinetics to CAR kinetics. (B) Antigen-independent (blue) and antigen-dependent (orange) intracellular cytokine production. Upper panel and bar graph show dual positive, TNFa+IFNg+ T cells at 18 hr time point, whereas lower panels show time course for single cytokine production of TNFa, IFNg, and IL-2. (C) Co-culture of CAR and OT-1 with left panel showing surviving OT-1s as proportion of OT-1s in co-culture Mock T cells. Right panel shows ratio of CARs:OT-1s over time, showing whether there is evidence of bi-directional fratricide.

However, there was a significant difference in the kinetics of killing with the α-Vβ5 CAR T cells killing over 90% of the target OT-I CD8 T cells within 24 hours, while the OVA-H2K^b^ CAR T cells took 48 hours to approach the same level of killing. There was a reduction in the CAR populations during this co-culture, demonstrating there is bi-directional fratricide from these interactions, but that the CARs ultimately prevail.

### pMHC-I CAR T cells are more effective than anti-Vβ CAR T cells at depleting polyclonal target T cells

We next looked to assess the efficacy the CAR constructs to kill target CD8 T cells *in vivo*. We sought a target T cell population that contained (a) a polyclonal TCR population more reflective of the variable TCR affinities that are most often found within autoimmune and other T cell-mediated diseases and (b) an immunodominant epitope. First, we utilized a well-characterized ovalbumin-expressing *Listeria monocytogenes* (LM-OVA) model in which C57Bl/6 mice are infected with LM-OVA, resulting in rapid generation of an effector CD8 T cell pool within the liver, of which a reproducible proportion are reactive to the immunodominant epitope from ovalbumin, SIINFEKL. Two days after infection, α-Vβ5 and OVA-H2K^b^ CARs were given to the mice to deplete the SIINFEKL-reactive T cell population. The mice were euthanized on day 10 and T cells isolated from the mouse livers. While *in vitro* data lead us to hypothesize that the α-Vβ5 CAR would be more effective at killing these target T cells due to its more rapid kinetics of activation and killing, the OVA-H2K^b^ CAR demonstrated greater depletion of the SIINFEKL-H2K^b^ tetramer-reactive T cells in this model. (Fig. 3a)

**Figure 3.**
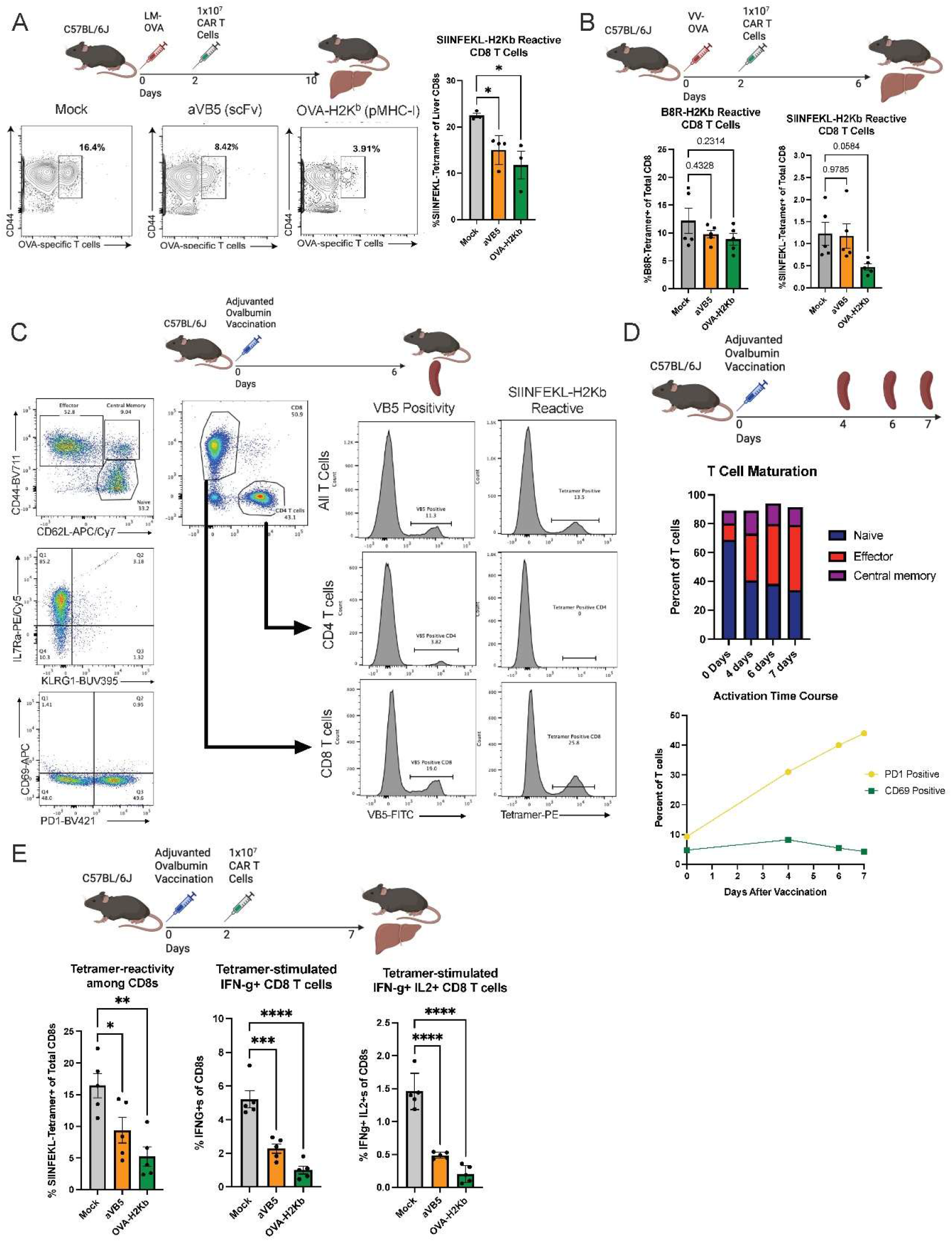
pMHC-I CARs are more effective than scFv CARs at killing polyclonal target CD8 T cells *in vivo*. (A) Experimental setup. B6 mice were infected with OVA-expression *Listeria monocytogenes*, given CAR T cells 2 days later, then euthanized at Day 10 to harvest livers and measure hepatic T cells response. Livers were macerated and enzymatically disassociated, then stained with tetramer and congenic makers and run on flow cytometry. (B) Experimental setup. B6 mice were infected with OVA-expression Vaccinia virus, given CAR T cells 2 days later, then euthanized at Day 6 to harvest livers and measure hepatic T cells response as before, with exception of use of two tetramers for SIINFEKL and B8R. (C) Adjuvanted vaccination with full length ovalbumin plus α-CD40 and poly(I:C) induces are repeatable polyclonal ovalbumin-reactive T cell population, detect in part here by SIINFEKL-reactive CD8 T cells.

To demonstrate the precision of the CARs for targeting specific epitope-reactive T cells, we utilized an infection model with vaccinia virus-encoding ovalbumin (VV-OVA). In this model, CD8 T cells respond to immunodominant peptides SIINFEKL from ovalbumin and TSYKFESV from vaccinia (the B8R peptide) as displayed by the MHC H-2K^b^. The OVA-H2K^b^ CARs decreased the number of SIINFEKL-reactive T cells without significant effect on the B8R-reactive T cells.

To further explore these findings, we next utilized an adjuvanted ovalbumin vaccination model.^10^ Ovalbumin plus anti-CD40 antibody and poly(I:C) are given via tail vein injection, which reliably generates a robust, activated effector CD8 T cell pool with high proportion of SIINFEKL-reactive T cells as demonstrated by the large proportion of CD62L^low^CD44^high^ and PD1+ T cells.(Fig. 3c) This adjuvanted-vaccine model generates memory precursor effector cells (MPECs), shown by the dominant KLRG1^low^IL-7Rα^high^ population. In this model, vaccination with full length ovalbumin generates a CD8-skewed population with increased proportion of Vβ5+ and SIINFEKL-reactivity. The generation of this robust effector T cell response begins early and peaks on day 7.(Fig. 3d)

We tested the ability of the α-Vβ5 and OVA-H2K^b^ CARs to deplete the SIINFEKL-reactive T cells by giving a dose of 1×10^7^ CAR T cells on day 2, followed by euthanasia and liver harvest on day 7. Once again, the OVA-H2K^b^ was more effective at depleting the SIINFEKL-reactive T cells. Additionally, fewer of the remaining CD8 T cells in this arm were able to produce IFNγ or both IFNγ+IL-2 in response to SIINFEKL tetramer.(Fig. 3e)

### Adoptive transfer of well-described adjuvanted ovalbumin-vaccinated T cells into RIP-mOVA reliably generates type 1 diabetes

Though there are induced models of T1D that rely on monoclonal T cell populations and spontaneous models of T1D that involve polyclonal T cell populations, there is not a polyclonal induced model of T1D. To address this gap in the field, we first tried direct vaccination of RIP-mOVA mice with ovalbumin plus aforementioned adjuvants.(Fig. 4a) However, this was unable to break tolerance in the RIP-mOVA mice, likely due to central tolerance mechanisms from ovalbumin expression in the thymus, preventing the generation of ovalbumin-reactive T cells. Thus, we gave the adjuvanted-ovalbumin vaccination to wild type mice and then adoptively transferred T cells into RIP-mOVA mice.(Fig. 4b) The ovalbumin-reactive T cells generated a robust hyperglycemic response that correlated to the amount of T cells transferred.

**Figure 4.**
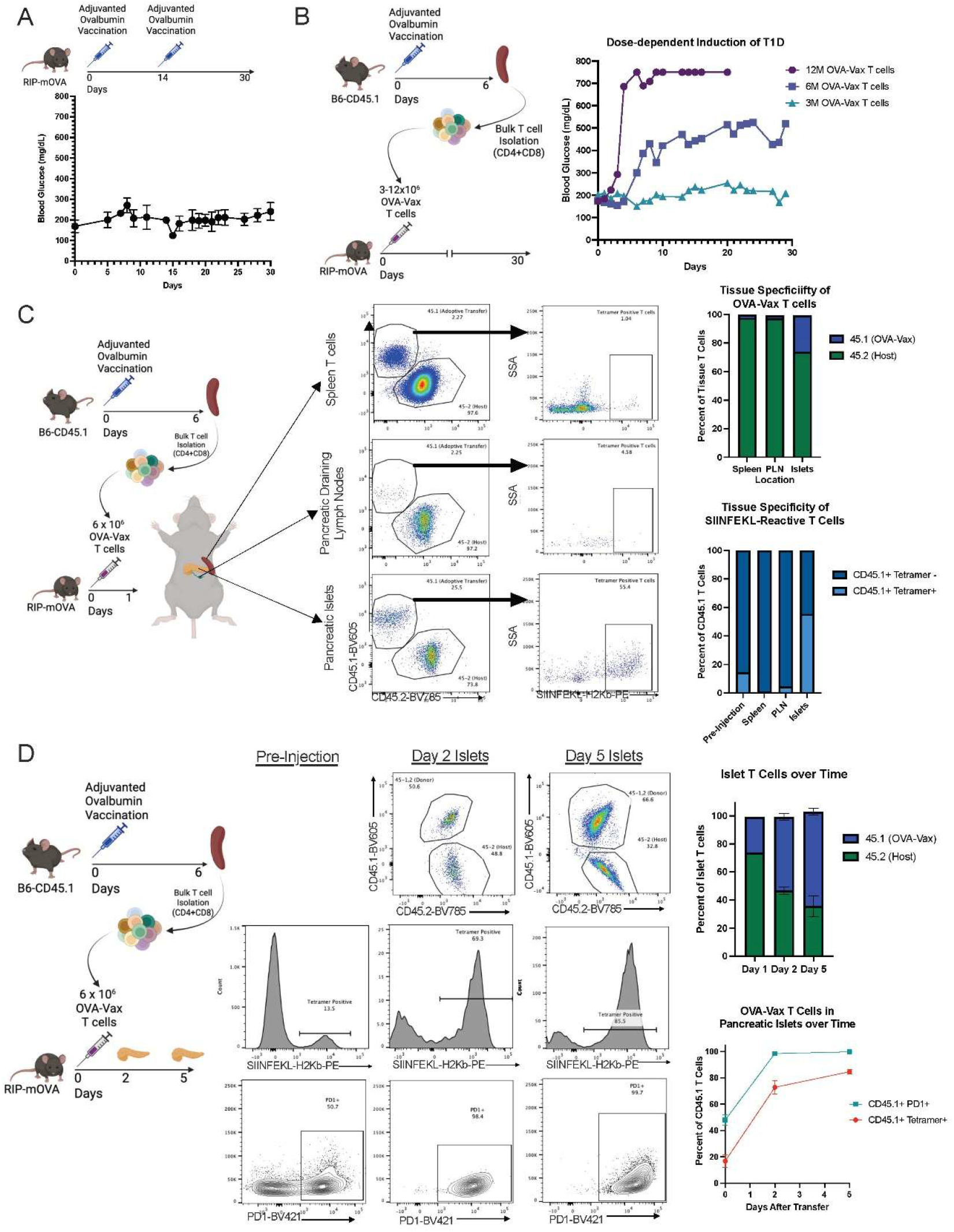
Development of novel polyclonal induced model of T1D in RIP-mOVA mice. (A) Experimental model of direct adjuvanted ovalbumin vaccination of RIP-mOVA mice with corresponding glucose curve from serial tail vein glucose checks. (B) Novel model of adjuvanted ovalbumin vaccination of PepBoy mice, followed by isolation of bulk splenic T cells to transfer into RIP-mOVA mice. Dose-finding experiment with 3, 6, and 12×10^6^ T cells given and RIP-mOVA mice monitored for development of hyperglycemia. (C) Experimental design to study organ penetration into spleen, pancreatic draining lymph nodes, and pancreatic islets by OVA-Vax T cells 1 day after adoptive transfer. Tetramer positivity is shown as percentage of adoptively transferred T cells in that organ. (D) Experimental design to study the activation of OVA-reactive T cells in pancreatic islets the first week after adoptive transfer, with *ex vivo* flow analysis showing proportion of OVA-Vax, SIINFEKL-reactive, and activated PD1^+^ T cells.

**Figure 5.**
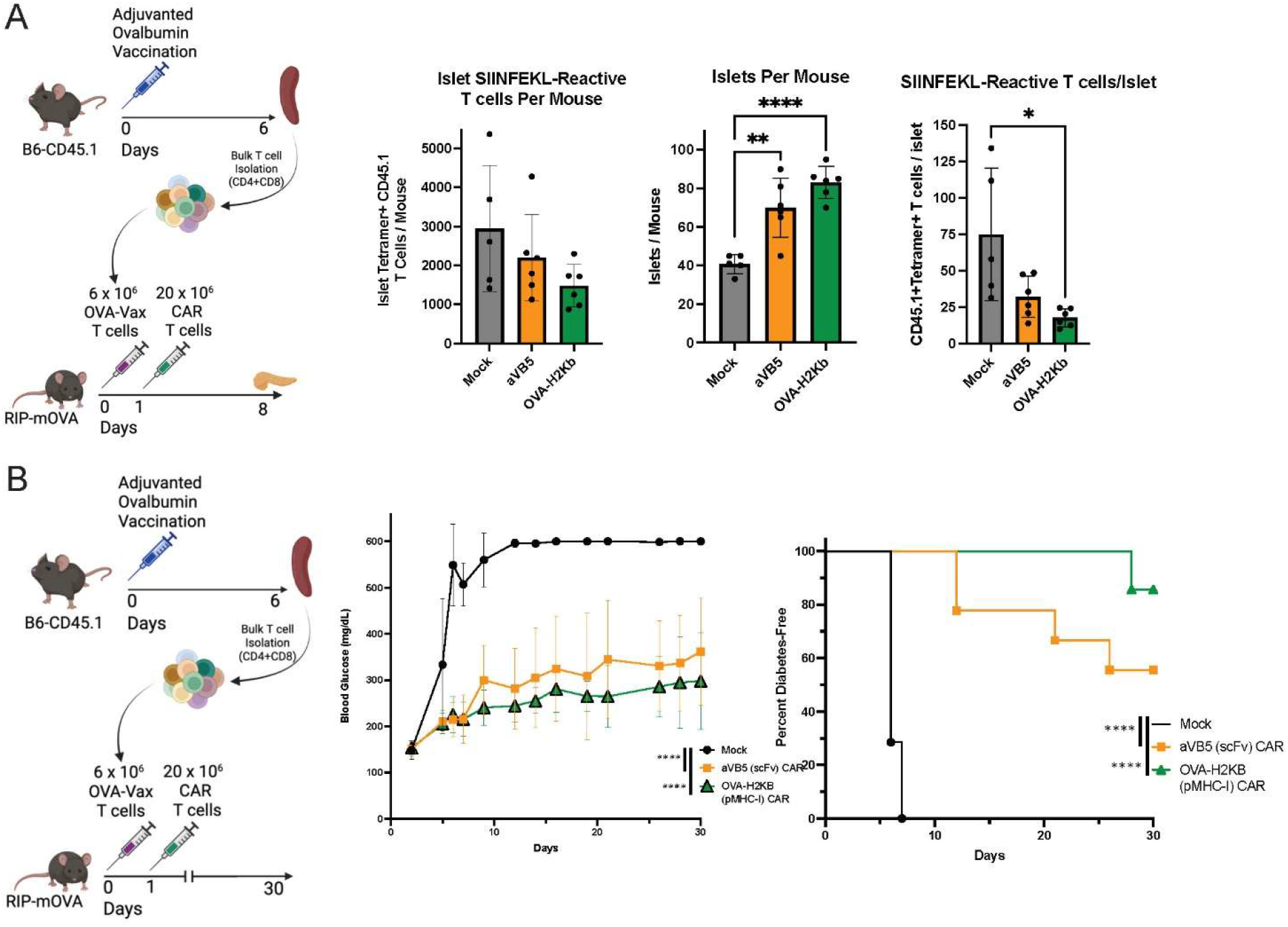
pMHC-I CARs deplete polyclonal autoreactive T cell population and prevent T1D. (A) Experimental model that builds on prior to add CAR T cells 24 hrs after adoptive transfer of diabetogenic T cells. Mice are euthanized at day 8 to isolate, count, and then disassociate islets and measure T cell populations. SIINFEKL-reactive T cells from the islets are shown per mouse, and then as a ratio to number of islets. (B) Experimental model that is same as before, but mice are monitored for the development of hyperglycemia and T1D, defined as 2 or more days in a row with a tail vein glucose reading over 300.

Upon transfer into RIP-mOVA mice, the adoptively transferred cells bypass the pancreatic draining lymph nodes and preferentially travel to and arrive in the pancreatic β-islets within 24 hrs of transfer, at which point 25% of total T cells within the β-islets are adoptively transferred are adoptively transferred and over 50% of these T cells are SIINFEKL-H2K^b^ tetramer binding. This islet T cell population rapidly expands and evolves, such that by day 5, 66% of the pancreatic β-islet T cells were adoptively transferred, of which 85% are SIINFEKL-H2K^b^ tetramer reactive and all are highly activated as measured by PD1^Hi^. This leads to destruction of the β-islets with subsequent hyperglycemia beginning at day 5-10.

### pMHC-I CAR T cells are more effective than anti-Vβ CAR T cells at preventing type 1 diabetes

Given that the pathogenic, ovalbumin-reactive T cells are present within the islets by day 1, we gave these mice high doses (15-20×10^6^) of the OVA-H2K^b^ and α-Vβ5 CAR T cells 24 hrs after adoptive transfer of ovalbumin-vaccinated T cells. One week later (day 8), we analyzed the pancreatic β-islets from these mice and show that the mice which received CAR T cells had more remaining islets and fewer SIINFEKL-H2K^b^ tetramer reactive T cells. While both CARs demonstrated protection of the islets from the ovalbumin-reactive T cells, the OVA-H2K^b^ showed greater killing of the SIINFEKL-reactive T cells. When these mice are monitored by glucose checks from tail vein bleeds for one month, all mice receiving Mock-transduced T cells developed type 1 diabetes (defined as two consecutive glucose readings over 300 mg/dL) by day 7 with glucose readings over 600 mg/dL by day 14. By contrast, the mean glucose readings went over 300 at day 15 for mice receiving the α-Vβ5 CAR T cells and did not cross this threshold in 30 days in mice receiving OVA-H2K^b^ CAR T cells. At day 30, 1/7 of mice receiving OVA-H2K^b^, 4/9 of mice receiving α-Vβ5, and 7/7 of mice receiving Mock T cells had become diabetic.

## Discussion

In this study, we compare the relative efficacy of a traditional scFv-based second generation CAR with a novel peptide-MHC Class I (pMHC-I) CAR in targeting and killing T cells via their TCR. Our *in vitro* studies demonstrate that both CAR formats are activated in the presence of target T cells. Despite also activating the target OT-I T cells, both CAR formats were able to withstand these cytotoxic effects and kill the OT-I T cells. The scFv CAR was able to activate and degranulate faster than the pMHC-I CARs, resulting in faster killing of *in vitro*. However, this benefit in kinetics did not translate into improved killing in our *in vivo* models utilizing polyclonal target T cells. Instead, the pMHC-I CARs were more effective in targeted killing of the ovalbumin-reactive T cells, likely due to increased precision leading to fewer targets to attack.

One concern that has been raised in previously published studies of peptide-MHC Class II CARs targeting pathogenic CD4 T cells is the difficulty that can arise in targeting a breadth of TCR affinities for epitopes of interest. To address these concerns withing our CD8-targeting CARs, we developed a variant of the RIP-mOVA model that utilizes adjuvanted vaccination of wild type mice to induce a polyclonal T cell population which is then transferred into RIP-mOVA mice where it reliably induces diabetes.

However, T cell driven autoimmune diseases often target multiple epitopes on the same antigen in a persistent and chronic manner. To more closely replicate this, we used full length ovalbumin vaccination of wild type mice adjuvanted with α-CD40 antibody and poly(I:C). This induces a Memory Precursor Effector Cell (MPEC) phenotype with a broad range of TCR cognate epitopes from ovalbumin which is capable of robust and persistent attack on the β-islets of the RIP-mOVA mice. Only 12-15% of these T cells are Vβ5 positive and/or bind SIINFEKL-H2K^b^ tetramer and thus able to be targeted by αVβ5 and OVA-H2Kb CARs, respectively. Despite this, these CARs reduce the number of SIINFEKL-reactive T cells within the β-islets and delay or prevent the onset of hyperglycemia in the mice. This demonstrates that depletion and control of a dominant antigenic epitope can control a much broader autoimmune response.

In the data presented here, both ovalbumin-reactive CD4 and CD8 T cells were given to the RIP-mOVA mouse. In our testing, this was necessary for the development of T1D in this model. Despite the presence of both classes of T cells, our CD8-targeting CARs were able to prevent or delay the induction of T1D. This is likely due in part to RIP-mOVA being a disease model that is largely dependent on pathogenic CD8 T cells, as compared to the NOD model of T1D which is more dependent on CD4 T cells. Regardless, this demonstrates the proof of concept that control of CD8 T cell-driven diseases is feasible through this approach even if there are other autoreactive T cell phenotypes present.

One weakness of this study is that the CAR T cells were not able to be detected long term. This is likely due to a combination of factors. First, this was done in an immune competent model, when many CAR studies require use of immunodeficient mice or immune conditioning regimens. We did not use this approach so as to more closely model clinical disease in our mouse models. Second, our CARs both used CD28 co-stimulatory domains and fully active CD3ζ internal signaling domains. This decision was made due to concerns for the risk of the CARs being killed upon binding to and activating the target CD8 T cells. In the future, we plan to optimize the CARs for persistence over early activation, through use of alternate co-stimulatory and signaling domains as well as rational modifications to the extracellular, TCR-binding domains.

## Methods

### Mouse Strains

B6.SJL-*Ptprc*^*a*^*Pepc*^*b*^/BoyJ (‘PepBoy’; strain 002014), Ins2-TFRC/OVA (‘RIP-mOVA’; strain 0054321) and C57BL/6J mice (‘B6’; strain 000664) mice were obtained from The Jackson Laboratory. Female mice were used for all experiments. All CAR T cells were generated from female donor mice at 6–10 weeks of age, and all hosts receiving adoptive transfers were female 6–12 weeks of age. All mice were bred and/or maintained in the animal facility at University of Colorado Anschutz Medical Campus. Mice were housed at up to five mice per cage with free access to chow (Inotiv, 2920X) and water. Statistical calculations for animal cohort sizes were based on conservative anticipated decreases in leukemia burden from 60% to 20%, with a standard deviation of 20%, a two-sided *α* of 0.05 and power of 0.80, with allowance of occasional unpredictable attrition during the study, resulting in a final cohort size of *n* = 5 biological replicates (mice) per group. The cohort sizes were generalized across studies based on the power analysis calculations specified above as well as our experience of the typical numbers of mice used in prior work. Mice were carefully monitored before experiments to ensure health before inclusion in experimental cohorts. For these reasons, we did not use additional mice beyond the numbers in our calculated sample sizes. Mice were housed at a temperature of 22 °C (±1 °C) at a humidity of 35%, with lighting 14 h on (0600–2000 h) and 10 h off and air changes at 10 to 15 per h. All experiments were performed in compliance with the study protocol approved by University of Colorado Anschutz Medical Campus Institutional Animal Care and Use Committee (protocol number 00751).

### Mouse DNA constructs

Basic construction of the mouse 1928z CAR was previously described.^11^ The mouse anti-Vβ5 scFv was Flag tagged to enable CAR detection, and all ITAMs in the CD3ζ domain were kept intact. A truncated human EGFR reporter protein was incorporated following a 2A skip sequence to provide an additional method for detection of CAR-transduced cells.^12^ The DNA was codon optimized, ordered from Thermo Fisher GeneArt and cloned into the MSCV-IRES-GFP backbone, a gift from T. Reya (Departments of Pharmacology and Medicine, University of California San Diego School of Medicine, La Jolla, CA, USA, Addgene plasmid 20672; http://n2t.net/addgene:20672; RRID: Addgene_20672) using XhoI and ClaI enzyme sites. The DNA sequences for the anti-Vβ5 scFv and OVA-H2K^b^ ectodomains were codon optimized, ordered from Thermo Fisher GeneArt and cloned into MSCV-IRES-GFP upstream of the IRES using EcoRI and XhoI enzyme sites.

### Cell lines and media

Mouse T cells and leukemia were cultured in complete mouse medium consisting of RPMI 1640 medium (Gibco) with 10% heat-inactivated fetal calf serum (Omega Bio), 1% nonessential amino acids (Gibco), 1% sodium pyruvate (Gibco), 1% penicillin/streptomycin (Gibco), 1% l-glutamine (Gibco), 1.5% HEPES buffer (Gibco) and 50 μM 2-mercaptoethanol (Sigma-Aldrich). All cell lines were routinely tested for myco-plasma (at least on an annual basis). Platinum E cells were purchased from Cell Biolabs and grown to create frozen stocks in Plat E medium containing DMEM, 10% heat-inactivated fetal calf serum (Omega Bio), 1% penicillin/streptomycin (Gibco) and 1% l-glutamine (Gibco) with 1 μg ml^−1^ puromycin and 10 μg ml^−1^ blasti-cidin for selection. Transfections were performed without penicillin/ streptomycin or selection antibiotics in the medium.

### Generation of retrovirus and lentivirus

For generation of mouse retrovirus, 1 × 10^6^ Platinum E cells (Cell Biolabs) were thawed and plated in a T75 flask and allowed to grow for 4 days. Cells were then split and re-seeded into T75 flasks at 12 x 10^6^ per flask and allowed to adhere overnight. The following day, the medium was changed to fresh Plat E medium, and transfection was performed with transfer plasmid DNA and Lipofectamine 3000 and p3000 reagents in basal OptiMEM medium (Gibco) according to the manufacturer’s protocol. Supernatant containing virus was collected 48 h later and spun at 2,000*g* to remove cellular particulate.

### Mouse CAR transduction

CAR transduction was performed as previously described.^13-15^ Briefly, spleens from 6-to 12-week-old donor mice were collected, and CD8^+^ T cells were isolated using an EasySep Mouse CD8^+^ T cell Isolation kit from STEMCell Technologies or bulk T cells were isolated using a Mouse CD3^+^ TCell Enrichment Column kit (R&D Biosciences, MTCC-25). On day 1, T cells were activated on anti-CD3/anti-CD28 Mouse T cell Activator Dynabeads (Gibco) at a 1:1 cell:bead ratio and cultured at 1 × 10^6^ cells per ml in complete mouse medium in the presence of recombinant human IL-2 (rhIL-2; 40 IU ml^−1^) and rhIL-7(10 ng ml^−1^) from R&D Systems. On days 2 and 3, retroviral supernatant was added to RetroNectin-coated (Takara Biosciences) six-well plates and spun at 2,000*g* at 32 °C for 2–3 h. The supernatant was then removed, and activated T cells were added to the wells at 1.0 ml per well. On day 4, beads were removed, and T cells were resuspended at 0.5 × 10^6^ cells per ml in fresh medium with cytokines. CAR transduction was determined after debeading by analyzing T cells by flow cytometry for a EGFR^+^ population plus surface expression of CAR via conjugated flow antibodies against FLAG or H-K2^b^ (or EGFR^+^ for control T cells).

### Vaccine and Adoptive Transfer model

The ovalbumin vaccine consists of 100 μg of whole ovalbumin protein (InvivoGen, vac-pova-100), 40 μg of anti-mouse CD40 (BioXCell, BE0016-2) and 40 μg of polyinosinic:polycytidylic acid (InvivoGen, tlr-5) per mouse resuspended to 200 μl total volume in PBS.^10^ For adoptive transfer models, bulk T cells were isolated from naive 6-to 8-week-old PepBoy mouse splenocytes using a Mouse CD3^+^ T Cell Enrichment Column kit (R&D Biosciences, MTCC-25). Tetramer staining was performed with PE-SIINFEKL-H2K^b^ tetramer to confirm presence of SIINFEKL-reactive T cells and adjust dosing as needed. Bulk T cells were then transferred into RIP-mOVA mice via tail vein injection. Mice were monitored for the development of hyperglycemia via tail vein blood draw and/or euthanized at pre-specified endpoints for islet isolation. This was performed by the Barbara Davis Center Islet Core. *Ex vivo* spleen, lymph node, and islet tissue underwent RBC lysis with ACK Lysing Buffer (ThermoFisher) and islets were further disassociated with collagenase prior to staining for flow cytometry.

### Flow cytometry

Flow cytometry analysis was performed using an LSR-Fortessa X-20 flow cytometer (BD Biosciences) and analyzed using FlowJo (BD Bio-sciences). Monoclonal antibodies used for staining are listed in the Reporting Summary. Intracellular flow cytometry staining was performed using the TrueNuclear Transcription Factor Buffer Set (Bio-Legend) for ex vivo staining of TFs, the Cytofix/Cytoperm Fixation/ Permeabilization kit (BD Biosciences) for intracellular cytokine staining.

### Mouse CD107a degranulation, intracellular cytokine staining, Ki67 and CellTrace dilution in vitro assays

In vitro assays were performed using a 1:1 effector:target cell ratio with 1 × 10^5^ of each cell type in a 96-well round-bottom plate, followed by analysis by flow cytometry at the indicated time points. Degranulation assays were performed by incubation for 6 h in the presence of 2 μM monensin and 2 μl of anti-CD107a. Intracellular cytokine staining was performed by incubation for 6 h with 1 μM monensin and 2.5 μM Brefeldin A added at 1 h in. All mouse in vitro assays were performed in complete mouse medium.

### Statistics

Statistical tests for all experiments except sequencing analyses were performed using GraphPad Prism v9.0 and v10.1.1 for MacIntosh (GraphPad Software). All data are represented as mean ± s.d. unless otherwise noted. Normality and equal variances of data were not formally tested; data distributions were assumed to be normal, and non-parametric or corrected tests were always used in cases where data were clearly outside of test assumptions. Data collection and analysis were not performed blind to the conditions of the experiments. No data points were excluded from analysis. Significance is denoted by \**P* < 0.05, \*\**P* < 0.01, \*\*\**P* < 0.001 and \*\*\*\**P* < 0.0001. Specific tests used along with technical, biological and experimental replicates in each dataset are indicated in the figure legends.

